# Sleep is required for neural network plasticity in the jellyfish *Cassiopea*

**DOI:** 10.1101/2023.05.04.538973

**Authors:** Michael J. Abrams, Lilian Zhang, Konnor von Emster, Brandon H. Lee, Hannah Zeigler, Tanya Jain, Ali Jafri, Zhiqin Chen, Richard M. Harland

## Abstract

Sleep in animals plays roles that appear specific to the brain, including synaptic homeostasis [1], neurotransmitter regulation [2], cellular repair [3], memory consolidation [4], and neural plasticity [5,6]. Would any of these functions of sleep be relevant to an animal without a brain? The upside-down jellyfish *Cassiopea xamachana*, like other cnidarians, lacks a centralized nervous system, yet the animal sleeps [7]. By tracking the propensity of the radially spaced ganglia to initiate muscle contractions over several days we determined how neural activity changes between sleep and wake in a decentralized nervous system. Ganglia-network sleep/ wake activity patterns range from being highly specialized to a few ganglia, to being completely unspecialized. Ganglia specialization also changes over time, indicating a high degree of plasticity in the neural network. The ganglia that lead activity can persist or switch between sleep/wake transitions, signifying a level of local control of the behavioral state in a decentralized nervous system. Following sleep deprivation, ganglia usage becomes far more sleep specialized, demonstrating reduced network plasticity. Together, these findings identify a novel behavioral control system that is decentralized and yet displays temporal specialization and centralization, and show a role for sleep in maintaining neural network plasticity, revealing a conserved function of sleep in this brain-less animal.

---

Though sleep is pervasive in animals, its fundamental roles, and the processes involved in generating the behavior, are poorly understood. An expanding set of model systems—fish, flies, and worms—have been used to address potential functions of sleep (Figure 1A). Although these species are members of different phyla, they all have a centralized nervous system (CNS), which receives information and coordinates the activity of the animal in a top-down fashion. Recently, sleep has been characterized in cnidarians [7,8], implying decentralized control of sleep, though there are varying degrees of local centralization within Cnidaria [9]. Three behavioral characteristics define a sleep state: quiescent periods, homeostatic regulation of quiescence, and latency to arousal (LTA) during quiescence [10]. In accordance, *Cassiopea* (Figure1B) displays quiescence, a slower pulse rate at night, which we can measure as an increase in the Inter-Pulse-Interval (IPI). If deprived of this nighttime quiescence they experience a rebound, a compensatory low activity period, the following day (homeostasis) [7]. During nighttime quiescence they also have a delayed response to stimuli (LTA) [7]. The presence of sleep behavior in this early branching metazoan provides the opportunity to investigate possible ancestral functions of sleep.

**Figure 1.**
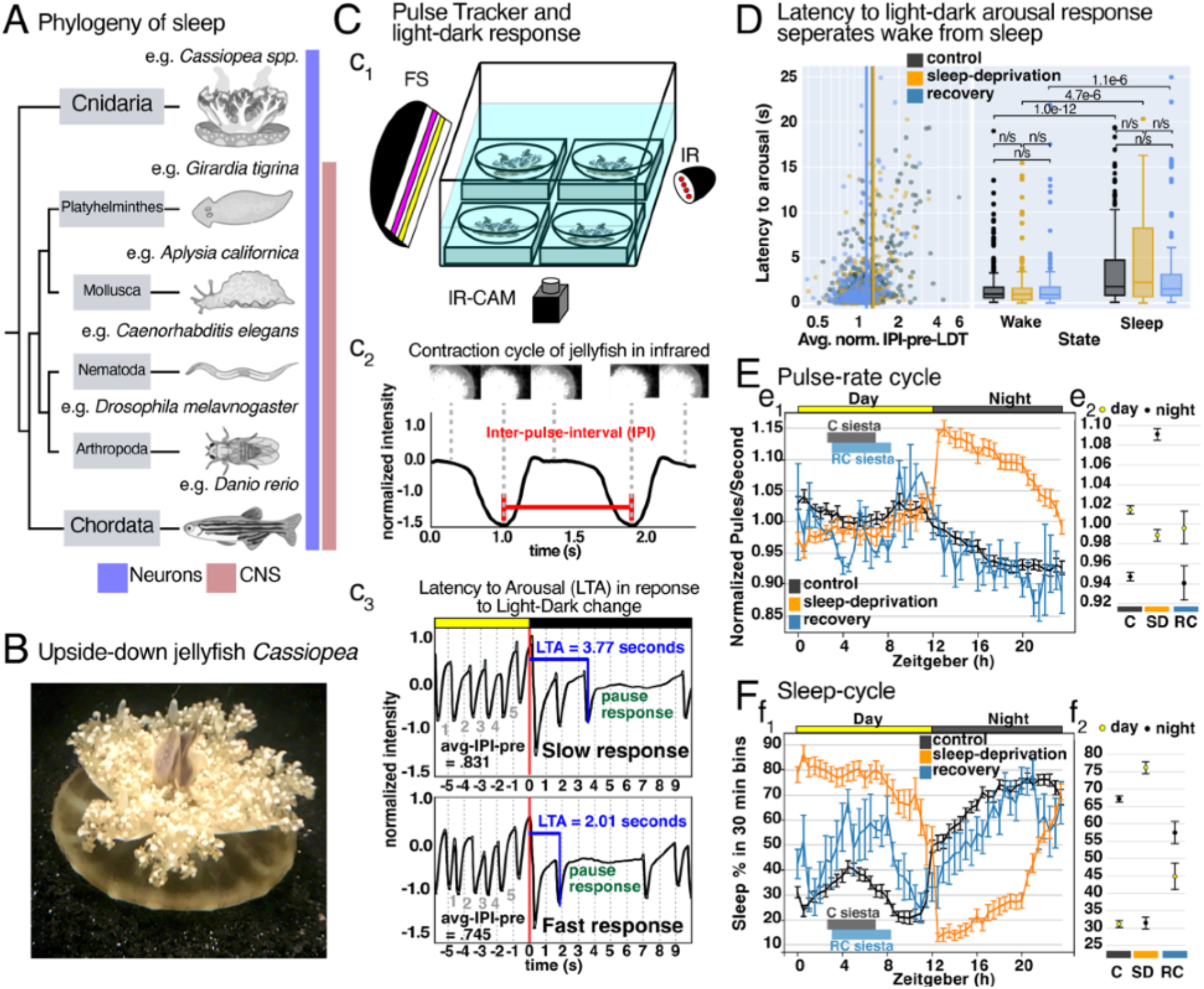
Phylogeny of sleep, *Cassiopea* pulse-tracking, and using Latency to Arousal (LTA) to detect sleep bouts in *Cassiopea*. **(A)** Phylogenetic tree highlighting the six phyla with described sleep states, adapted from [16], and the lack of a CNS in *Cassiopea*. **(B)** *Cassiopea* in its natural upside-down orientation. **(C)** For Pulse Tracker recordings, **(c**_**1**_) jellies are placed in square watch glass dishes, and recorded in infrared (IR) at 15 frames per second (CAM). **(c**_**2**_) The average pixel intensity is calculated every frame within a region of interest (example image series shown above the pulse trace), and normalized to remove background light fluctuations. The frame with least intensity occurs when the jelly is most contracted. The time between contractions, the Inter-Pulse-Interval (IPI), is shown. **(c**_**3**_) Two example light-dark responses are shown, with the IPIs-pre and -post light-dark change. Five IPIs pre-light to dark change are shown (grey) for each trace, and their respective 5-IPI average (avg-IPI-pre). The time, LTA (blue), until a pause response (green) is detected; in this case the pause response is defined by an IPI ≥ 2X avg-IPI-pre (1.5X to 5X show similar trends, Supplemental Figure 1D). **(D)** A cycle of 5 min in light, 1 min in dark, for 10 cycles was complete on 14 animals, for six consecutive days for control, two consecutive days for sleep-deprived, and three consecutive days for recovery was completed. **Left panel**, the average normalized activity pre light-to-dark change for each jelly is plotted against their corresponding latency to arousal. A median threshold, vertical lines, was calculated (see Methods) for control (black), sleep-deprived (gold), and recovery (blue). **Right panel**, box plots quantifying the LTA of below threshold (wake-like) and above threshold (sleep-like) responses. p-values calculated using one-way ANOVA, followed by Wilcox rank sum test. **(E-F)** Pulse-tracker information for control (C), sleep-deprived (SD), and recovery (RC) animals. n = 25 control, 19 SD, 6 RC. Pulse-rate **(e**_**1**_) and % sleep **(f**_**1**_) binned every 30 min, for night vs day, error bars are SEM. Pulse-rate **(e**_**2**_) and % sleep **(f**_**2**_) binned by day (yellow) or night (black), error bars are 95% CI.

## Sleep-bout characterization in *Cassiopea*

To determine possible functions of sleep in *Cassiopea* we first needed to establish criteria to distinguish wake from sleep. Previously, *Cassiopea* was shown to take longer to arouse to a mechanical stimulus at night than during the day, by allowing the the animal to fall from a stationary position in the water [7]. In order to associate specific neural patterns with sleep or wake, here we defined a behavioral activity threshold to segregate the two states. We characterized discrete periods of sleep using a new latency-to-arousal (LTA) tests [10-14] in two steps: 1) a non-aroused activity level was determined, and then 2) the response time to an arousing stimulus was measured. *Cassiopea* have a strong pause response when lights are turned off, perhaps akin to the hyper-arousal freeze response of other animals [15]. Previously, “Pulse Tracker” (Figure 1c1&c2), which finds the average pixel intensity within a region of interest, was used to find the pixel intensity local minima that coincide with muscle contractions [7]. We used Pulse Tracker to quantify the average of 5 IPIs-pre light change (aver-age-IPI-pre, Figure 1c3), in both light-to-dark and dark-to-light transitions, to gain non-aroused light and dark behavior (Supplemental Figure 1A), completing step 1. Some animals pulse faster than others, so to determine the non-aroused behavior among *Cassiopea*, it is necessary to normalize activity of individual jellyfish by their mean activity (Supplemental Figure 1B). Then, for step 2, we measured the latency to the pause induced by the light-to-dark test. (Figure 1c3). We used the median of the normalized average-IPIs-pre LDT as the activity threshold (Supplemental Figure 1C) to separate wake-like from sleep-like pulsing activity (Figure 1D, left panel).

Using this approach, we cycled animals through 5 min of light and 1 min of dark, and found a significant increase in LTA for animals that are pulsing slower, in control (p-value 1e-12), sleep deprived (p-value 4.7e-6), and recovery (p-value 1.1e-6) conditions (Figure 1D, right panel, p-values generated from one-way ANOVA, and post-hoc Wilcox ranked sum test). As part of this test, a response threshold is required to determine the moment when an animal is aroused (paused), we tested several response thresholds from 1.5X to 5X the average of 5 IPIs-pre light-to-dark test, which all yielded similar results [Supplemental Figure 1D]. Further, similar results were obtained using mechanical perturbations to confirm the the separation of wake from sleep both day and night (Supplemental Figure 1E-F). We also tested dark acclimation length, and found that SD animals given 5 min of dark have a large increase in LTA compared to control-sleeping animals (Supplemental Figure G, p-value 1.8e-8), implying a deeper sleep state in SD animals when given greater time to dark-acclimate. Together, these experiments have defined a useful behavioral activity threshold to determine when *Cassiopea* are asleep or awake.

We tracked *Cassiopea* sleep patterns to see if they resemble those of other animals by recording *Cassiopea* pulsing for several consecutive days and nights in control, sleep-deprived, and recovery conditions (Figure 1E, Supplemental Figure 1H). A clear cycle of pulsing activity was observed: control animals progressively decreasing pulse-rate at night, sleep-deprived animals have a lower daytime pulse-rate and drastic increase in night-time pulse-rate, and recovery animals have a low daytime and an even lower nighttime pulse rate (Figure 1E). To identify sleep from wake, we found the median threshold IPI (the median of all light- and dark-IPIs) for each animal every 24-hour period. Control *Cassiopea* sleep 31.1±1.0% the day and 67.1±1.0% of the night (Figure 1H). Not-surprisingly, animals experiencing nighttime light pulses sleep little at night, 31.4±1.7% (Figure 1H), but exhibit a strong sleep rebound (compensatory low-activity period) the following day, sleeping 76.1±1.7%. Finally, after two nights of SD, recovering animals display an intermediate level of sleep both day and night (Figure 1H), sleeping 44.8±3.8% of the day and 57.4±3.2% of the night. Interestingly, both control and recovery animals experience a midday siesta, which is common among animals [34-36], but recovering animals undergo a larger and more continuous siesta (Figure 1F). These patterns share similarities with the sleep of other animals [17-19], and is further evidence of a conserved state.

## Tracking muscle contraction initiation sites acts as a proxy for ganglionic pacemaker activity

*Cassiopea* has a decentralized nervous system comprised of radially spaced (Figure 2A), morphologically identical rhopalia (Figure 2B) which are light and balance sensing organs. Each rhopalium contains a ganglion that is capable of controlling the behavior of the entire animal by initiating muscle contractions [20-23] that generate the pulsing behavior that allowed us to characterize the animal’s sleep state [7]. In *Cassiopea*, the network of pacemaker-containing ganglia controls the pulse rate, so presumably it must also facilitate the decrease in average pulse rate during sleep. There are fundamental similarities between pacemaker networks in different jellyfish, including electrical coupling of ganglia [20-23]. Modeling work in the box jellyfish *Tripedalia cystophora* indicates that ganglionic pacemakers with the highest activity initiate contractions and hyperpolarize their ganglionic competitors [29]; therefore, tracking contraction-initiating ganglia would detect the pacemakers with highest activity. To confirm earlier work showing that an electrical signal originates from *Cassiopea* ganglia [22], rather than from a non-discrete nerve ring as observed in distantly related jellyfish (Hydromedusae) [30], we recorded field potentials from rhopalia dissected away from the muscle. We recorded compound action potentials from these isolated rhopalia (Figure 2c1-2, Supplemental Figure 2A), but no action potentials in tissue that did not contain ganglia. As with other jellyfish, the impulses that drive pulsing behavior of *Cassiopea* start in ganglionic pacemakers and arrive first to proximal muscle fibers; these impulses then propagate outward throughout the muscle, as in *Aurelia aurita* [31]. Therefore, initiation sites of muscle contractions act as a proxy for ganglionic pacemaker activity.

**Figure 2.**
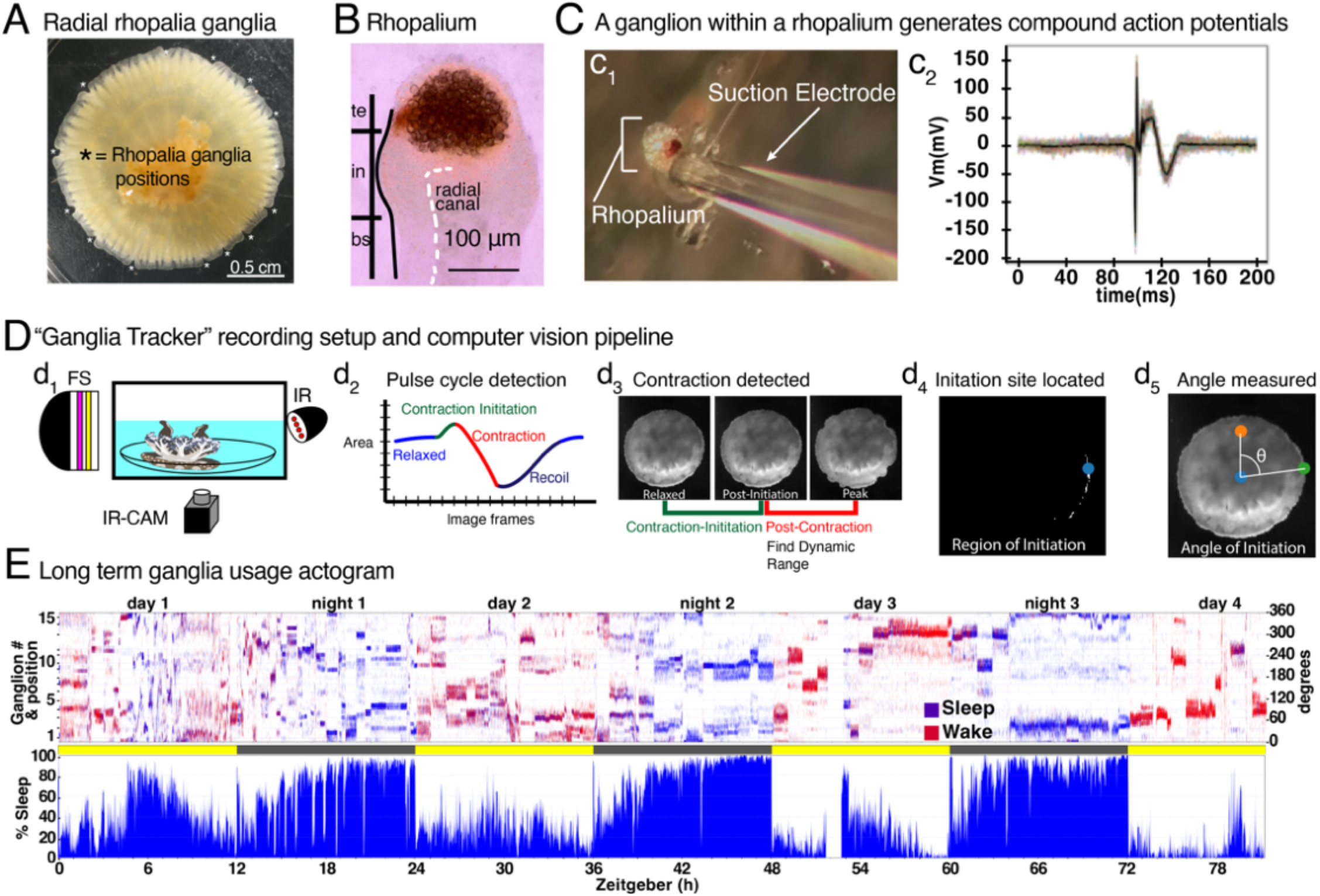
Electrophysiology supported optical method gives longterm high temporal and spatial resolution pacemaker activity. **(A)** Oral view of *Cassiopea*, asterisks indicate the radial spacing of the rhopalia. **(B)** Rhopalium, oral view, with the three labeled segments, te, terminal, in, intermediate, bs, basal, the radial canal. In the terminal segment the dark brown lithocyst is visible, and the light brown melanin can also be seen from the spot ocelli on the aboral side. **(C)** Ganglionic action potentials; **(c**_**1**_) rhopalium mounted on suction electrode, **(c**_**2**_) and 22 sorted and aligned compound action potentials. **D)** Ganglia tracker setup and computer vision pipeline. **(d**_**1**_) Cassiopea cannot escape, or maneuver such that their bell margins are obscured, and are maintained in full-spectrum (FS) light, on a 12:12 cycle, fed every third day, and are recorded in IR with a 120 fps camera (CAM). **(d**_**2**_) The image analysis program detects the area of the jellyfish every frame, and detects the period immediately pre-and post-contraction, **(d**_**3**_) and then the contraction periods are further analyzed for the position of the jellyfish and finds specific moments from each pulse that are used **(d**_**4**_) downstream to detect location of muscle contraction initiation, blue dot. **(d**_**5**_) Finally, the angle (theta) of the the initiation site relative to the 0 degree point (red dot) is determined. **(E)** An example actogram showing four contiguous days and nights is shown. y-axis is ganglion number and position, with 0-degree at the bottom and 360-degrees at the top. Ticks are pulses, and colored based on if they are above (sleep, blue) or below (wake, red) the median-IPI their associated 24-hour day-night cycle. Below the actogram is a bar that indicates lighting, under which is the average sleep (blue) in 1 min bins. At the bottom is the Zeitgeber hour.

## Ganglia show specialized temporal hierarchies that exchange homeostatically and lead the animal through sleep-wake transitions

To study how the distributed ganglia in a decentralized neural architecture might control pulsing during sleep and wake, or dark and light, or during and after sleep deprivation, we developed a “Ganglia Tracker” to locate the region of muscle-contraction initiation using high speed video, and by proxy, the ganglion that initiated the contraction (Figure 2D, Figure 2B-E Supplemental). This optical method gives us high temporal and spatial resolution to assess longterm pacemaker activity, and exploits the relatively simple neural architecture of *Cassiopea* to address the most basal and fundamental functions of sleep.

We used Ganglia Tracker to assess ganglion pacemaker activity over time, and to assign wake (red) and sleep (blue), we used a threshold median IPI for every 24-hour day-night period for each animal. We plotted actograms for individual jellyfish over several days, with the state (sleep vs. wake) and position (polar coordinates) of each contraction initiation (Figure 2E). Actograms do not appear as random noise making it immediately obvious that, (1) certain ganglia lead contractions for long periods, others lead only rarely, and 2) ganglionic leadership is not a constant and can change between days as well as within a single day.

Ganglia usage displays three distinct patterns during transitions between behavioral states: maintain leadership activity, turn on leadership activity, or turn off leadership (Figure 3A). For examples, at zeitgeber (ZT) 0, ganglia 1, 2 and 10 maintain their sleep activity in the dark as the animal wakes in the light, while ganglia 4 and 12 turn on activity, (Figure 3a1 cyan asterisk, and Figure 3b1); at ZT 5, ganglion 14, which had been previously active during sleep and wake, turns off, and ganglion 4 takes on most sleep activity while ganglia 6-7 lead most wake activity (Figure 3a1 fuchsia asterisk and Figure 3b2); at ZT 9, ganglia 1, 3 and 9 maintain leadership as the animal transitions between wake and sleep (Figure 3a1 green asterisk, and Figure 3b3). Actograms show that ganglia network is locally controlling the behavioral state of the animal as it transitions between wake and sleep states using ganglia in both specialized and unspecialized ways.

**Figure 3.**
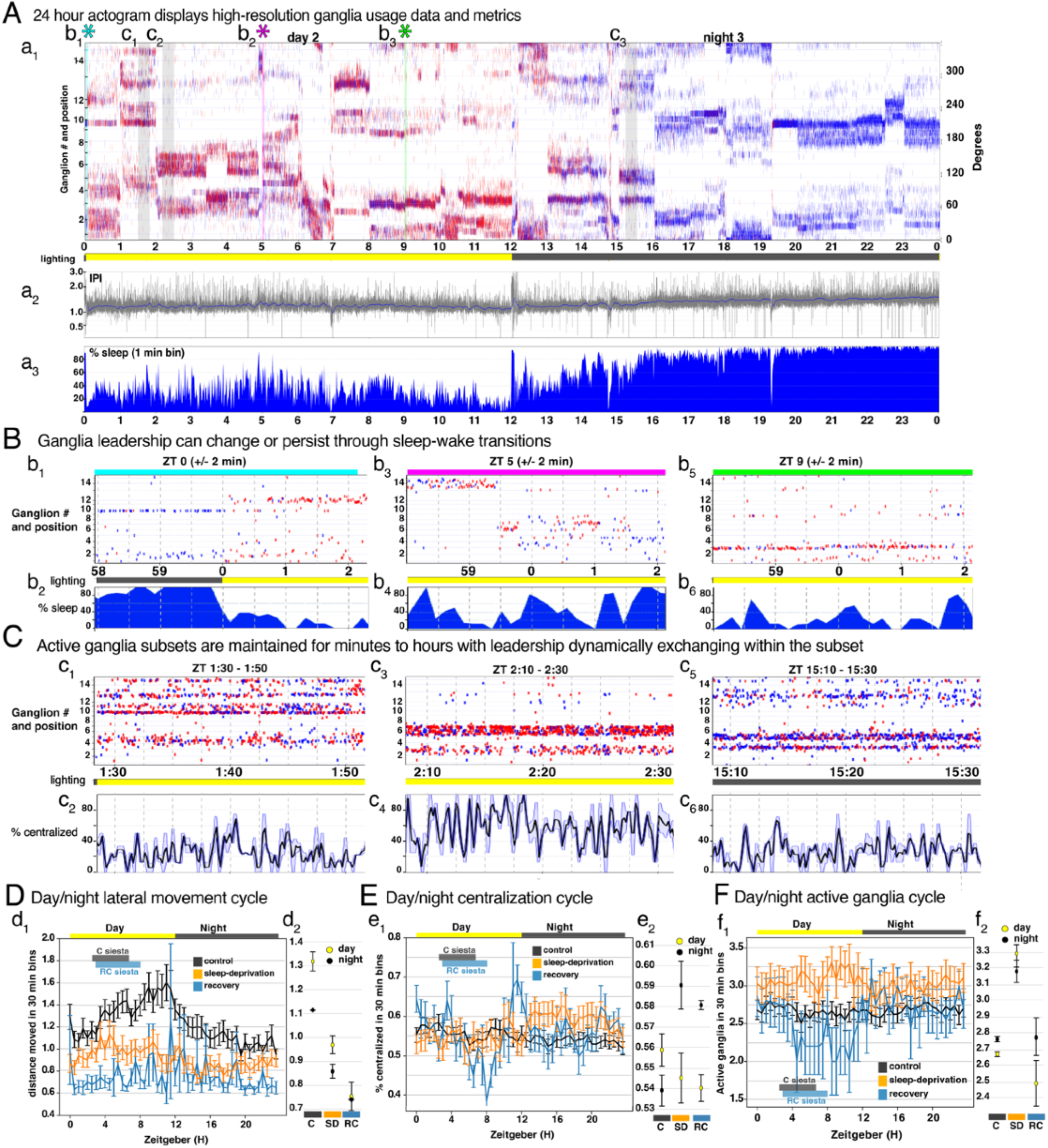
Sleep/wake ganglia activity tracking reveals long- and short-term hierarchies and possible specialization. **(A)** 24-hour actogram of day 2 and night 3 magnified from Figure 2E. **(a**_**1**_) actogram data, under which are indicators for day (yellow) and night (dark grey), **(a**_**2**_) IPI, and **(a**_**3**_) percent sleep in 1min bins). **(B)** Ganglia leadership during sleep/wake transitions, actogram data from 4 minutes, with % sleep (20 second bins). **(b**_**1**,**3**,**6**_) Examples of ganglia maintaining, turning on or off leadership during **(b**_**2**,**4**,**6**_) sleep/wake transitions. **(C)** Actogram data for 20 minutes and % centralized (20 second rolling average (light blue ± SEM). **(c**_**1**,**3**,**5**_) Examples of stable ganglia leadership subsets, **(c**_**2**,**4**,**6**_) that show rapid fluctuations in centralization. **(D-F)** Control (black), sleep-deprivation (gold), recovery (blue), n = 14 control, n = 12 SD, n=5 RC. **(d**_**1**,_ **e**_**1**,_ **f**_**1**_) Average lateral movement (pixels), Average network centralization, Average number of active (above 10% of total activity per bin), in 30 min bins, error bars are SEM or, **(d**_**2**,_ **e**_**2**,_ **f**_**2**_) in 12 hour bins, error bars are 95% CI.

## *Cassiopea* appear to have nervous system specialization with dynamic ganglionic interactions that are affected by sleep deprivation and recovery

We developed a centralization metric to assess how the network is dynamically used over time, calculated by determining the frequency of consecutive pulses originating from the same ganglion—displayed as a 20-second rolling average (Figure 3c2,c4,c6) A centralization value of 0.5 equates to contraction leadership alternating between two ganglia, each with equal leadership frequency. Centralization levels can fluctuate rapidly, but trends appear towards lower (Figure 3c2,c6) and higher (Figure 3c4) centralization. This network metrics indicates that leadership switches between a small subset of ganglia, with the most active initiators intermittently losing leadership to the next most active competitor.

In animals with a CNS, different areas of the brain are specialized for specific functions [32]— for example, some areas of hypothalamus promote sleep while others promote feeding [33]—and this specialization is stable throughout the animal’s life. Our data indicates that despite lacking an anatomically-defined CNS, *Cassiopea* too have nervous system specialization. But in contrast with a CNS, this spatial-temporal specialization is plastic within each animal.

We used Ganglia Tracker to study how neural network activity patterns change during sleep deprivation and recovery. Lateral movement (periods when an animal is moving laterally in the recording setup), centralization, and active ganglia (those that initiate at least 10% of all contractions in 30 minutes, giving us the size of the active network), all appear to cycle levels between day and night, and all are impacted by sleep deprivation (Figure 3D-F). There are several notable features. First, for lateral movement, the control daytime movement appears higher than the night, but there is a stall in the progressive increase through the day during the siesta; sleep deprivation reduces lateral movement, and recovering animals move least (Figure 3D). Second, for centralization, control animals show higher centralization during the day than at night, with a local minimum during the midday siesta, which is more severe in recovering animals; also, compared to controls, SD animals show much higher centralization at night, indicating a possible correlation between nighttime arousal and increased centralization (Figure 3E). Finally, the number of active ganglia shows a slight decrease in controls, but a more major decrease in recovery animals, during the midday siesta (Figure 3F), while the SD animals have a larger active network both day and night. The recovery conditions during the siesta is perhaps the most revealing, in which a smaller active network may be allowing the larger set of inactive ganglia to recover, while simultaneously minimizing the additional neural work to the smallest functional network. Together, sleep behavior in *Cassiopea* involves multiple cycling attributes, indicating that there are substantial changes in the nervous system between wake and sleep that are also affected by SD and recovery.

## Wake-active ganglia change more between light-cycles than sleep-active ganglia

To assess ganglia specialization and homeostasis we used actogram data to compare the top-most active ganglia between 12-hour light cycles, and allowed us to determine how much the dominant ganglia change their sleep or wake activity between cycles. From this example actogram (Figure 4A), we identified the pulses that occur during sleep (Figure 4B) and wake (Figure 4C), and calculated the top two-, five-, and eight-most active sleep or wake ganglia in each 12-hour cycle (top-two, green; top-five, green, and magenta; top-eight, green, magenta, and grey). We then measured the change in their activity between 12-hour cycles; in this example the difference between night and day among the night-most-active ganglia (Figure 4d1-d3). The sum of the absolute differences yields the total activity change between one 12-hour cycle; in this case, there is a greater difference in activity in wake-ganglia compared to sleep-ganglia (Figure 4d4). We also saw that much of the difference, 51% of wake and 33% of sleep, came from the two most active ganglia. Are these trends consistent between control animals, and what is the affect of sleep deprivation?

**Figure 4.**
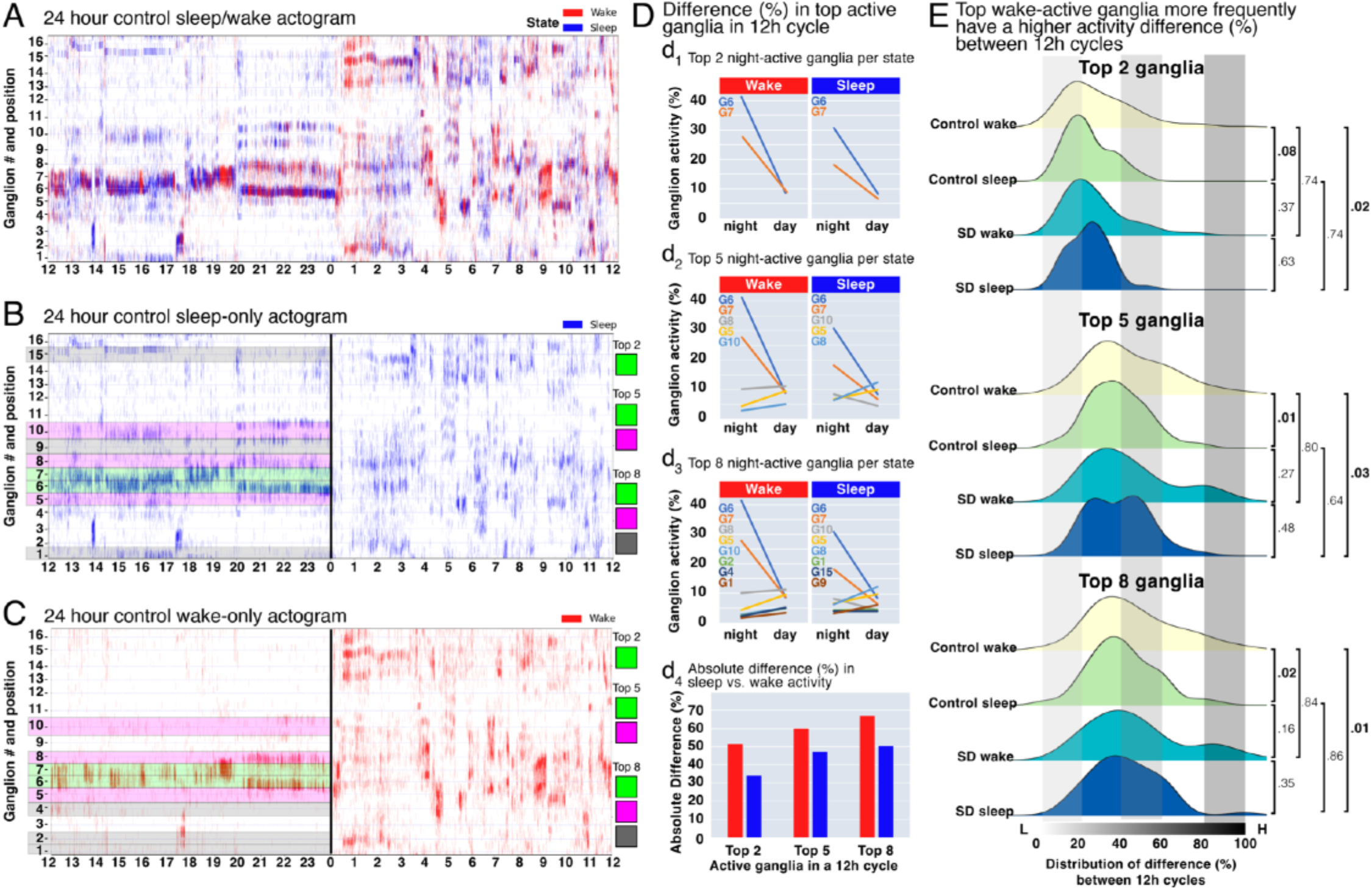
Wake-active ganglia change activity more between 12-hour light cycles than sleep-active ganglia. **(A)** A 24 hour actogram. **(B-C)** The pulses during sleep **(B)**, and wake **(C)**, with the top 2 most active ganglia highlighted in green, the next three (3-5) most active ganglia in magenta, and the next three (6-8) most active ganglia in grey. **(D)** The ganglia activity from night to day in the top 2 **(d**_**1**_), top 5 **(d**_**2**_), and top 8 **(d**_**3**_) ganglia. Each ganglion is labeled G1-16. **(d**_**4**_) The total absolute difference in ganglia activity in the most active ganglia in this example. **(E)** Ridge plots compare the distributions of ganglia activity differences between 12 hour periods, between control wake, control sleep, SD wake, and SD sleep, for the top 2 **(e**_**1**_), top 5 **(e**_**2**_), and top 8 **(e**_**3**_) most active ganglia. Plots display the differences generated from 39 control animals with 162 transitions, and 18 SD animals with 57 transitions. p-values generated by Kolmogorov-Smirnov tests.

We compared the absolute difference in sleep- and wake-ganglia percent activity systematically between 162 control, and 57 SD, 12-hour light cycles, to determine if there are overall trends. A value of 0.5 means that half of the ganglia activity from one 12-hour cycle has switched to different ganglia for the next 12-hour cycle. Between all conditions, the top-two, -five, and -eight, have average differences of, 20±1.6%, 35±1.8% and 45±2.0%, respectively (range indicated is the 95% CI). Notably, there is a shift towards a higher difference in the top-most wake-active ganglia compared to the top-most sleep-active ganglia (Figure 4e1, control sleep vs control wake, p-value .08; Figure 4e2, control sleep vs control wake p-value .01; Figure 4e3, control sleep vs control wake p-value .02), implying there may be an added cost to wake activity. At the same time, sleep-active ganglia appear more longterm activity specialize, similar to the impact of sleep deprivation. This approach is focused on the activity of specific ganglia in 12-hour cycles, so next we assessed at finer time resolution how the entire active ganglia network specializes.

## Sleep deprivation impacts neural network specialization and plasticity

To determine the extent of neural network specialization in *Cassiopea*, 24 hours of actogram data from control (Figure 5a1), SD (Figure 5b1), and recovery (Figure 5c1) animals were processed using principle component analysis [37], a technique that transforms a dataset to give each point a coordinate that describe its variability in fewer dimensions. Each point in a PCA is the linear transformation of 1-minute bins of actogram data, using the activity of each ganglion as its features (Supplemental Figure 3A). Then, k-means clustering [38], measuring the intra-cluster sum of squared errors from the centroid, is used to group bins into clusters (Figure 5a2, b2, c2). The actogram can be recolored to show the cluster identity for each contraction to gain confidence visually in the clustering (Figure 5a3, b3, c3). Finally, the average sleep activity of the bin can be overlaid on the attogram PCA to ascertain the percentage of bins in each cluster that occurred during sleep or wake, allowing us to calculate the sleep specialization of each cluster is shown (Figure 5a4,b4,c4). The same analysis was performed to assess night or day specialization. We tested how many clusters tend to exist in actogram data using the approximate elbow of k-means inertia plot (Supplemental Figure 3B), and tested how well the clustering maps back to the actogram data (Supplemental Figure 3C), and found that 8 clusters reasonably distinguishes actogram patterns.

**Figure 5.**
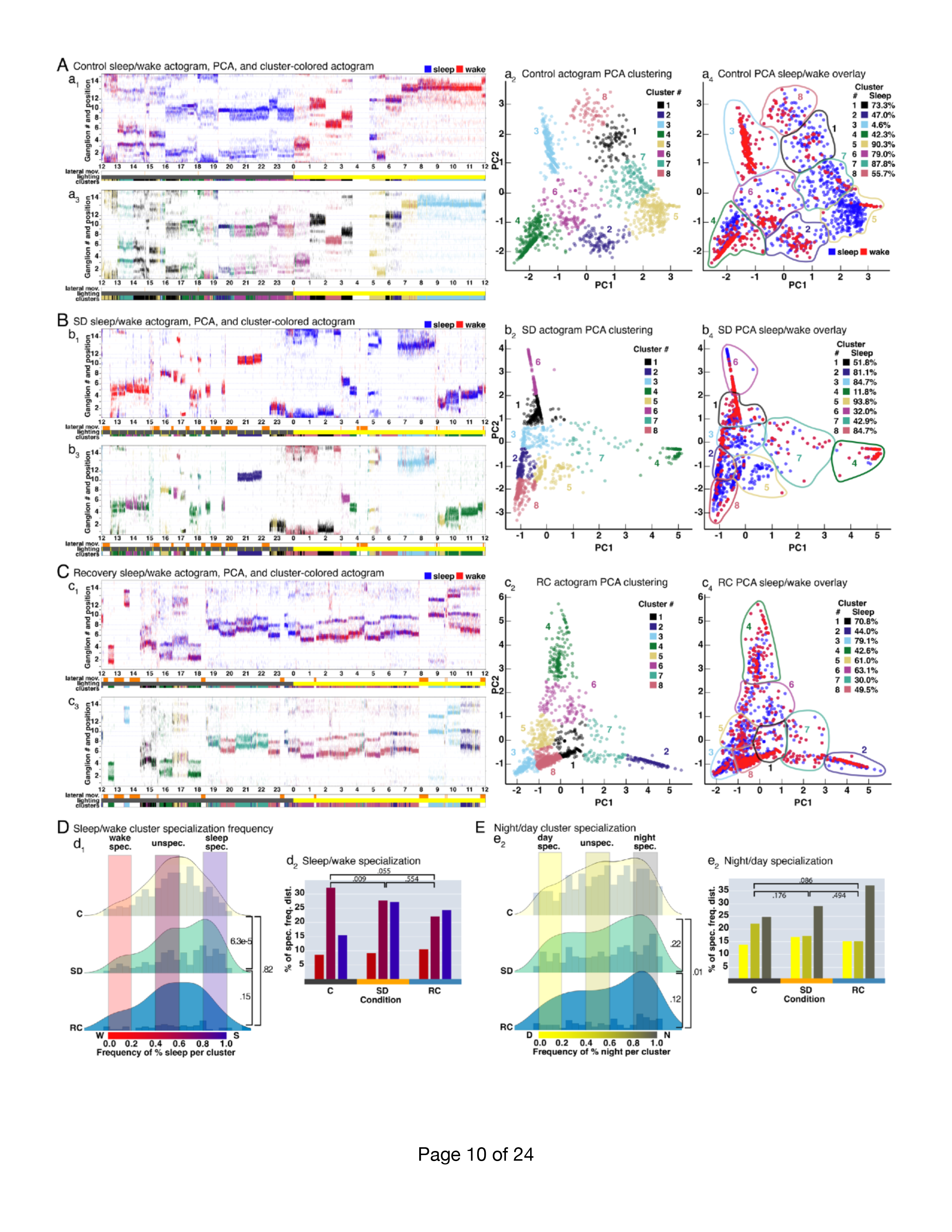
PCA of ganglia usage reveals the extent of sleep/wake and night/day specialization in *Cassiopea*, and that sleep deprivation increases sleep specialization. **(A-C)** Actograms from a single animal from **(a**_**1**_) Control, **(b**_**1**_) SD, and **(c**_**1**_) RC recordings 24-hour recordings, and the associated PCAs **(a**_**2**_) Control, **(b**_**2**_) SD, and **(c**_**2**_) RC, using ganglia activity 1-min bins, and k-means clustering. Clusters in all animals had a silhouette score of 0.1 or higher and had more than 5 bins each. Colors and numbers indicate cluster. The actograms were then re-colored using the cluster color associated with each 1-min bin for **(a**_**3**_) Control, **(b**_**3**_) SD, and **(c**_**3**_) RC. The sleep/wake state of the bin is colored based on its average activity for **(a**_**4**_) Control, **(b**_**4**_) SD, and **(c**_**4**_) RC. **(D-E)** PCA sleep/ wake and night/day specialization analysis of n = 14 control animals with 385 clusters, n = 14 SD animals with 223 clusters, and n=5 RC animals and 54 clusters. Distributions **(d**_**1**_,**e**_**1**_) of the specialization % for each cluster, separated by condition, visualized using matplotlib Gaussian kernel density estimator. Y-axis is normalized density, and X-axis is the frequency of **(d**_**1**_) % sleep, or **(e**_**2**_) % night, per cluster; 1 is maximum sleep, **(d**_**1**_) or night **(e**_**1**_) specialized, 0 is maximum wake **(d**_**1**_) or **(e**_**1**_) day specialized, and 0.5 is unspecialized. p-value determined by Kolmogorov-Smirnov test. Categorical counts, normalized to total count, for each state and condition, given as the percent of the cluster specialization frequency distribution. Sleep **(d**_**2**_) or night **(e**_**2**_) specialized (0.8 - 1.0), **(d**_**2, e2**_) unspecialized (0.4-0.6), and wake **(d**_**2**_) or day **(e**_**2**_) specialized (0.0 - 0.2). p-values determined by 2×3 chi-square.

We consider network specialization to be the tendency for specific combinations of ganglia (clusters) to drive muscle contractions, and whether certain clusters are present at different frequencies associated with behavior state (sleep/wake) or light conditions (night/day). For example, cluster 3 (cyan) identified using PCA (Figure 5a2, a3), exists primarily when ganglia 13 and 14 are highly active, during ZT 8-12. These ganglia are active together only 4.6% during sleep, indicating that it is a highly wake specialized cluster. In contrast, if we look earlier that day, at cluster 5, primarily active from ZT5-6, 7-8 (the midday siesta) consisting of ganglia 2, 4, and 12, 13, 15; these ganglia are 90.3% sleep-active, so it is highly sleep specialized. While some ganglia change their activity, overall specialization patterns persist over time (Supplemental Figure 4A-D).

To understand how cluster specialization applies broadly, among many individuals, or between conditions, we plot distributions with the specialization density obtained through a Gaussian kernel density estimator. The cluster specialization frequency, from 14 control, 14 SD, and 5 recovery animals, was normalized to the number of clusters per condition, making the distribution shapes directly comparable for sleep/wake (Figure 5D) and night/day (Figure 5E) specialization. The control sleep/wake distribution shows highest specialization density near the center (Figure 5d1,d2), in the unspecialized region, indicating that patterns present during sleep are often also present during wake—evidence of a high level of network plasticity, or a lack of specialization in active ganglia. However, we do see a full spectrum of specialization, showing that some clusters are still active only during sleep or wake. To see any degree of cluster specialization in this systemic way is intriguing, as the null hypothesis would be that in a decentralized nervous system there should not be any specialization. We used a permutation test, which randomly assigned bin sleep/wake or night/day specialization, to test this null hypothesis, and found the sleep/wake and night/day cluster specialization is significantly different from the null distribution generated by permuting bin labels (Supplemental Figure 3D). We also confirmed the robustness of clustering by using a drop out test, which randomly dropped two bins from each cluster (Supplemental Figure 3D). With the recent characterization of functional modules in *Clytia* [39] and distinct neural population controlling various types of of muscle contractions in *Hydra* [40], it seems likely that an unappreciated level of specialization occurs in Cnidaria.

Remarkably, sleep deprived animals become far more sleep specialized than controls (Figure 5d1, p-value 6.3e-5, Kolmogorov-Smirnov test; Supplemental Figure 4E), indicating clusters in sleep deprived animals are far more often active only during sleep. While sleep specialization increases, the frequency of SD unspecialized clusters decreases, without particularly affecting wake specialization, whether we consider specialization the more radical top or bottom 20% of the spectrum (Figure 5d2), or more liberal top or bottom 33% (Supplemental Figure 4E). We interpret this increased sleep specialization as a decrease in plasticity compared to control animals. Importantly, the shift in the distribution towards sleep specialization observed in SD animals is driven by daytime rebound sleep. Perhaps the deeper sleep state associated with dark-acclimated rebound sleep we characterized using LTA (Supplemental Figure 1G) involves a higher degree of specialization. In recovery periods, the distribution appears to return to a distribution similar to control (Figure 5d1), though there is still more sleep specialization compared to controls (Figure 5d2), indicating a level of reversibility, and a pressure to return to a more plastic neural state. The PCA and cluster specialization analysis supports a role for sleep in maintaining normal neural network specialization and plasticity, that is severely impacted by sleep deprivation.

Conversely, the night/day specialization distributions appear similar to each other, all three have rather flat distributions, implying there is a high degree of both night and day specialization, though there is a shift towards night specialization (Figure 5e1). Control and SD are not significantly different from each other, but recovery is further night specialized when we consider the top or bottom 20% of them spectrum as specialized (Figure 5e2), though this pattern is not significant if the top or bottom 33% is considered specialized (Supplemental Figure 4G). Our finding again is not impacted by the drop out test, and night/day specialization is significantly different from a null distribution generated by permuting labels (Supplemental Figure 3D). Therefore, again we see neural network specialization, but these distributions are significantly different than the sleep/wake specialization (Supplemental Figure 3D), implying we are measuring two distinct types of specialization occurring simultaneously in *Cassiopea*.

## A role for sleep in the nervous system of *Cassiopea*

There are two major theories for how global brain states arise to generate behaviors like sleep. Either specialized regions control the switch between wakefulness and sleep (a top-down mechanism) [41]; or neural networks have an emergent bias towards certain global states that are affected by local regulatory circuits (a bottom-up mechanism) [42]. From work in humans and model animal systems there is a growing appreciation that sleep is regulated globally, regionally, and locally by intersecting mechanisms [43]. While it is clear that sleep across animals plays many roles, perhaps it arose initially in a simpler form that is preserved in *Cassiopea* sleep.

In this study, we have characterized how *Cassiopea* ganglia are used in spontaneously occurring long-term hierarchies (of active and inactive, sleep vs. wake, and light vs. dark) that exchange overtime. These properties show a novel pattern in a previously uncharacterized form of nervous system, one that is anatomically decentralized and yet spatiotemporally centralized. There may be costs of leadership and neurological activity that might build up over time that necessitate or underpin the exchange—which could connect to other possible roles for sleep including mitigating cellular stress and replenishing key macromolecules [44]. Along this line, it will be interesting to determine if there are correlations between ganglia activity and molecular markers of stress, proliferation and cell death.

The ganglia exchange could be regulated, it will be important to assess what neural circuits control pacemaker activity and facilitate ganglionic competition, and how those circuits integrate a history of activity and environmental or state-based cues to generate ganglia hierarchies. As *Cassiopea* express many of the core neurological circuit components, there may be conservation in pacemaker regulation —cholinergic, glutamatergic, or GABAergic neurons have been implicated in regulating pacemaker activity in *Hydra* [45], the crab, *Cancer borealis* [46], zebrafish [47], and in rodents [48]. Also, *Cassiopea* can experience periods of extreme decentralization (Figure 3c1,c2 at ZT1:10, Figure 3c5, c6 at ZT15:10), and perhaps there are ways to induce this phenomena for us to study in detail. Disorganized pacemaker activity in *Cassiopea* may share traits with similar occurrences in other animals [49,50], giving us a new model to study this phenomenon.

It may also be that the network is constantly optimizing for the most efficient activity-maintaining network, and that the specialization we observe arrises spontaneously. Perhaps we are observing something similar to neural-network simulations that spontaneously form brain-like functional specialization when given specific tasks to achieve [51]. As all ganglia are capable of initiating contractions, we also wonder what the non-contraction initiating (inactive) ganglia are doing—perhaps they are resting, or are specialized for some other task.

Our finding that sleep-deprivation drives a shift towards sleep specialization of the ganglia network implies that sleep promotes the unspecialized state. Could an unspecialized ganglion state be favorable? It is interesting to consider the benefits of a more plastic network as it is difficult to imagine that it would facilitate learning or memory in *Cassiopea*. However, *Cassiopea* live in an environment with predators, intermittent prey availability, and breeding cycles [52], so perhaps the unspecialized system is better primed to respond to a changing environment. The plasticity could allow the animal to rest some ganglia without affecting behavior much, which may be similar to how some birds and ocean mammals have semi-hemispheric sleep [53], allowing half of the brain to sleep while the other maintains necessary activity, or another phenomenon, local sleep, that has been recorded in mammals, in which brain regions appear asleep even while the animal remains conscious [54].

## Author Contributions

M.J.A. conceived the project. M.J.A. and R.M.H oversaw the project. M.J.A., L.Z., H.Z., and A.J. conducted experiments. M.J.A., L.Z., K.V.E., B.L., H.Z., T.J., A.J., Z.C. carried out data analysis. M.J.A conceptualized, designed, and built experimental setups. K.V.E., L.Z.., and M.J.A wrote image-processing and data-analysis scripts. M.J.A wrote the paper with input from R.M.H.

## Acknowledgments

We thank Aki Ohdera and Lea Goentoro from Caltech, CA; and Victoria Sharp from Pennsylvania State University, PA, for generously supplying *Cassiopea* medusa and polyps. We thank Aki Ohdera and David Raizen for critical reading of the manuscript. We thank Marta Truchado Garcia and our Harland Lab colleagues for their stalwart counsel and technical guidance. We thank our compatriots in the International Cassiopea Workshop, CassiopeaBase, and the Key Largo Marine bResearch Laboratory, for input and discussion. We thank Milad Shafaei for his insight that helped form the Light-dark-test. We thank Dan Costanza for his early work in developing Ganglia Tracker. We thank Noah Olsman and Daniel Eaton for critical advice. We thank Nicole King for early support, and Richard Kramer for electrophysiology advice and instrumentation. We thank the incredible UC Berkeley undergrads who worked on Cassiopea behavior, Matthew Lim, Mayra Arellano, Tessa Auer, Shuangyue Li, Sofia Telfer, and Mira Patel; and Cassiopea data analysis, Kevin Lu, Edmond Chang, Haley Mozier, and William Ling. Thank you to the UC Berkeley Undergraduate Research Apprenticeship Program. This work was supported by the Miller Institute at University of California, Berkeley, and the C.H. Li Distinguished Professor fund.

## Materials and Methods

### Dye injections, oral arm trimming, and ganglion amputation

Medusa were placed in ice cold 800uM menthol for a few moments for immobilization. To ensure consistent orientation, each jellyfish received three dye injections. The dye was composed of india ink dried on 10uM glass beads (Sigma-Aldrich 440345), which was then powderized and resuspended in 2% PEG, dissolved in DI water. Medusa were placed in ice cold 800uM menthol for a few moments for immobilization, then three injection were made by hand with a large gauge (21G x 1 1/2) needle, to avoid clogging, one day prior to behavioral recordings, and generally persisted for several days. Oral arms were also trimmed back using small scissors (FST 14058-11) so they did not overhang the edge of the bell. Jellyfish are imaged using an IPhone camera at the time of dye injection, in 60mm petri dishes.

### Ganglia Tracker

#### Recording setup and approach

This behavioral recording setup consisted of 20 L clear hard plastic containers with their bottoms replaced with 1/4 inch thick clear acrylic. Three and four inch diameter watch glass dishes were placed on the acrylic with silicone grease, and several small pieces of clear silicone tubing were cut longwise so they clamp down on the glass rim, creating a bumper that prevented the jellies from escaping without preventing water flow. Pulaco PL-188 aquarium heaters kept the system temperature controlled at 84°F. Filtered water was constantly pumped into the containers and flows out of a short adjustable stand pipe. An automatic top-off system maintained the salinity of the recording setup between 28 and 32 parts per thousand (ppt) artificial sweater (Instant Ocean).

ACA1300-200UM Basler cameras with 10X Computer macro lens with BP850 IR filter were used. Cameras were placed about 1.5 feet beneath the ganglia tracker recording setup. The cameras connected to a Matlab program that recorded mp4 videos at 120 frames per second (fps). The lights used for the behavioral recording jellyfish were fixtures with two full spectrum T5 and two actinic T5 bulbs. The lights were on a 12 hour cycle from light to dark, where lights came on at 7 AM and turned off at 7 PM. SD experiments utilize bright LED lamps attached to a cycling timer that turns on 5min every 25min. Experiments followed the recording schedule shown in Supplemental Figure 1H).

#### Analysis Pipeline

##### Initializations

Due to the duration of recordings and size of the video files, behavioral analysis is divided between local computers and Savio, U.C. Berkeley’s High Performance Computing Cluster. The processing pipeline starts with an initialization step on local computers using a 30 second video segment representative of stationary pulsing behavior in subsequent full recordings. The purpose of these initialization steps is twofold: to assess the quality of the recording and to set parameters for downstream processing. Steps are then iterated in parallel over the full length recording period on Savio to obtain the angle of the initiating ganglia.

Recordings must be manually checked for video quality and pulse profile. By running Pre-Init.py on the local computer, the user will have generated a 3600-file image stack of the chosen 30 second chunk and the PreInitializationDF.csv, a data frame containing the number of frames, Savio file path, and frames per second of each recording. This data frame provides the basis for the information necessary to process recordings in parallel on Savio. To account for recordings truncated due to video corruption or pauses for feeding, the number of missing frames is added to the truncated file to ensure an accurate timeline is produced post-processing.

After the image stack and data frame is generated, the user may then run the *Initialization_Main*.*py* script on the 30 second chunk of recording. Following thresholding to convert grayscale images from IR recordings to binary images, the largest binary pixel area is utilized to identify the jelly area. Through this, a pulse profile of the jelly is generated with a regular behavioral pattern consisting of an initiation, peak, trough, and relaxed phase. Identification of these specific frames requires the user to manually set the optimal numConsecutiveDrops based on the pulse profile, which will denote the number of frames the binary jelly area must consecutively decrease to be labeled a trough. The manually set peak2InflectionDiff and peak2TroughDiff will respectively use the difference in frames from the peak to inflection point and the trough to peak to mark the inflection point and peak. To aid in processing efficiency, a postPeakRefractoryPeriod will also be set by the user, which will denote the number of frames in the refractory period to skip in downstream processing. Taking the binary difference of relaxed and trough images allows for identification of the centroid as the center of the largest area once noise is removed. The initiation site is determined by thresholding the difference between the image post-inflection point and the relaxed image. This threshold is chosen from a pre-set range of 0.1 to 0.5 based on the lowest standard deviation when comparing initiation angles calculated from the program to angles measured by eye. To continue with downstream processing, the standard deviation must be below 25 degrees.

##### Video Processing (Savio)

Following initialization, video chunks and the PostInitializationDF.csv containing all initialization parameters are transferred to Savio. *FFMPEG*.*sh* is then run to convert all .mp4s to .jpg image stacks in parallel, distributed over 24 overlay containers. This file system serves to decrease the load of write and delete steps on Savio’s HPC system and reduce image conversion time to an average of three hours for a three day recording.

Upon completion of *FFMPEG*.*sh, videoProcessing_job*.*sh* runs the processing pipeline using initialized parameters. Video processing on Savio uses the same methods as mentioned above in Initialization, iterated in parallel over the full recording. However, as the 30 second video chunk in initialization steps is of a stationary animal, lateral movement is accounted for by tracking the centroid position. The centroid is re-calculated every pulse. Upon a centroid difference greater than 20 pixels (as set in the initialization steps), the jellyfish will be considered in movement and angle data will not be obtained. Following a period of 7200 frames with a centroid at the same location, the jellyfish will once again be considered stationary and subsequently re-initialized to select new initiation, peak, trough, and relaxed frames. Due to changes in lighting that may occur with movement, the difference between these new frames will be used as new threshold and normalization images to obtain the initiation angle. The image following the movement phase will be saved to allow for orientation of the rhopalium based on dye markings post processing. Rhopalia positions are measured in Fiji, and the 0 degree mark is assigned to a dye mark that persisted through the entire experiment.

##### Post-Processing

Once processing on Savio is completed, angle data and orientation frames are produced. The angle to orient images following movement will be manually assigned by the user. Based on this, a complex DataFrame can be produced, which extracts data including the Zeitgeber time, closest rhopalia to the initiator, distance moved, and interpulse interval (IPI). Pulses with IPIs greater than the median IPI within 24 hours are labeled “sleep” and those less than the median are “wake”.

##### DeepLabCut

Using DeepLabCut [55], a ResNet-50 convolutional neural network was trained on an initial dataset of 24 hours of recording, with seventeen labels which were used to estimate the location of rhopalia. Since DeepLabCut is often used to track the different body parts on an animal, it was difficult for the program to locate and differentiate rhopalia. Instead, labels were initially placed on random points on the circumference of the jellyfish, which it would track throughout the whole duration of the video. After this processing, the positions of the rhopalia were measured by-hand and compared to these labels in order to offset them to the correct positions. By combining the values of the rhopalia positions with the angles of initiating ganglia, we were able to track ganglia activity levels for long periods of time, without needing by-eye reorientation after every moment segment.

#### Pulse Tracker

##### Recording setup and approach

**Light-dark-tests**, jellyfish were placed in 4in x 4in x 4in acrylic boxes filled with enough seawater to cover the jellyfish (approximately 1.5 inch of water height) in at 28-32 ppt artificial sweater. Four square watch glasses of 1 and ⅝ inch (Carolina Biological Supply) were placed in each box, with each watch glass holding one jellyfish for a maximum of four jellyfish per container. White netting was placed over the jellyfish to prevent to ensure that the jellyfish remained within their watch glass. Acrylic boxes were placed in a Pyrex dish filled with distilled water and with three Pulaco PL-188 heaters set to 84°F. This system resulted in a heated water bath for the jellyfish. Lighting was the same as with Ganglia tracker. The same camera and lens was used, but the Matlab program recorded mp4 videos at 15fps. The lights were on a 5 min digital timer, to cycle on and off, for 10 cycles, and an additional infrared light is on the timer cycle which is used to indicate when the lights turn on and off.

##### Pulse Tacker Light-Dark-Test processing

**To extract the inter-pulse-interval (IPI) of jellyfish**, we first put recorded videos through the program ffmpeg_parallel.sh, which used FFMPEG to convert multiple videos into image stacks simultaneously at a rate of 15 fps. Next, We ran Pulse Tracker Analysis Software as was done previously [7], but with some adjustments for infrared recordings. Briefly, the first image of the chosen timeframe is displayed to allow the user to select a rectangular region of interest surrounding each jellyfish. Inside of this region, the software measures average pixel intensity change over time—when the jellyfish is relaxed it has more light pixels and when it is contracted there are fewer. Maximum intensity indicates a relaxed jellyfish and minimum pixel intensity indicates full bell contraction. Then, plots are generated with the normalized pixel intensity change/time. When the trace passes a chosen negative threshold a pulse is denoted, shown with a red dot in. Graphs where the average pixel intensity change was not significant enough to allow for clear marking of pulses do occur, generally due to poor lighting conditions or if the animal has turned on its side, and these periods cannot be used for analysis. The output of this program has the time at which a pulse occurred and the IPI between the current and previous pulse.

##### Pulse Tacker Light-Dark-Test analysis

The exact frame at which a light change occurred is determined by eye, using the cycling infrared light mentioned above. A five minute period between light changes allows animals to acclimate before the next transition (Supplemental Figure 1A). The IPI information for every jellyfish is then imported into a python analysis and plotting script, which extracts the last 5 IPIs pre-lights-off and pre-lights-on, for each cycle. The average of each 5-IPI set is calculated generating the light- and dark-average IPIs (Supplemental Figure 1B, left panel in black). Each is then divided by their respective **normalization factor** (the median of the combination of all light-to-dark and dark-to-light average IPIs for that animal) for a **dimensionless normalized IPI** (Supplemental Figure 1B, right panel in green). A threshold is required to distinguish sleep from wake. We applied a **median threshold** (the combined normalized average of light- and dark-IPIs pre-LDT, Supplemental Figure 1C) for baseline, SD, and RC animals. Finally, to measure the LTA we find the **response time** (time from the light-dark switch until pause response). We tested various thresholds for defining a pause, and all showed segregation of sleep from wake (Supplemental Figure 1D). We placed 14 animals into the Pulse Tracker recoding setup, and cycled them through 5 min of light and 1 min of dark, 10 times, starting at zeitgeber (ZT) ∼6, for 6 consecutive control days, two consecutive sleep-deprivation days, and three consecutive recovery days. Animals were sleep deprived (SD) using 5 min pulses of light every 25 min, for each of the two consecutive nights. The median average-IPI-pre for control (black line), SD (gold line), and recovery (blue line) (Figure 1D, left panel) were used to threshold sleep from wake in their respective conditions (Figure 1D, right panel). To determine the impact of greater dark acclimation, we repeated the experiment on 14 more animals, this time using a cycle of 5 min of light and 5 min of dark, and collected two consecutive control days, and two consecutive sleep-deprivation days (Supplemental Figure 1G).

##### Drop test video recordings and analysis

**Drop-tests** require more space, and 3×2 array of clear cylinders with a taught brine-shrimp screen were placed in a 20 L clear plastic tank. The array had arms that rested on small boxes that keep the cylinder array about 2 inches from the top of the water. To drop the array, the small boxes that rest on the lip of the plastic tank were quickly pulled out, and the array falls and stops when the array arms land on the plastic tank rim. Drop-test recordings are done from above, with IR lights placed around the bottom of the tank. The IPI-pre and IPI-post drop, in both the light and dark, are assessed by eye from the video recordings. The IPIs of all pre-drop test are averaged, generating a **normalization factor** by which all IPIs are the divided. A **median threshold** (on normalized pre-drop IPIs in both light and dark), sorts wake-like from sleep-like normalized post-drop IPIs (Supplemental Figure 1E-F).

#### Actogram analysis

##### PCA Pre-processing

Clustering analysis was performed in Python, with modules from scikit-learn and using complex DataFrames generated through CassiopeaProcessor data analysis as described above. From these complex DataFrames, the data was cleaned to remove outliers of distance moved greater than 50 pixels and rows without identifiable initiating rhopalia. 24-hour periods of each jellyfish were then extracted for analysis, to reflect shifting day-to- day variability in pulsing behavior.

##### DataFrame generation

A dataframe at a granularity of every 10 minutes was then generated, using features including the fraction of usage of each rhopalia, the median interpulse interval, and the distance in pixels that the centroid has moved. The fraction of usage of each rhopalia was generated through the sum of contractions initiated by a given rhopalia, normalized to the total number of contractions within the 10 minute period. This fraction of usage was then thresholded at a 10% activity threshold. Data was centered and normalized prior to analysis.

##### PCA and k-means clustering

This data frame was then analyzed through Principal Components Analysis (PCA). Clusters were assigned through *k*-means clustering, using the inertia (intra-cluster sum of squared errors from the centroid) to choose the optimal *k* clusters for clustering PCA results. 10 minute bins whose median IPI was greater than the median IPI of the 24 hour period were assigned as “sleep-like” periods, and bins whose median IPI was lower than that of the 24 hour period were assigned “wake-like”. Bins that occur during the night or day are also assigned.

##### Cluster Activity Distribution

To measure patterns of behavior in aggregate, we then quantified the percentage of sleep-like vs wake-like 10-min bins were within a given cluster. The distribution of sleep-like cluster percentages was visualized as violin plots using the matplotlib Gaussian kernel estimator.

#### Electrophysiology

##### Assembly and analysis

To record from the whole ganglia, suction electrodes were made by pulling a 1 mm glass capillary on a capillary puller and breaking tips to a clean opening. Electrode size was compared to rhopalium to ensure adequate suction. Jellyfish (approximately 3 cm in diameter) were amputated along the radii to produce a strip of ganglia. Strip was placed oral-side up in a clean petri dish containing sterile filtered artificial seawater (SFASW). SFASW was then removed until water level was just above the rhopalia and the tissue lies flat against the dish surface. Suction electrode was filled with the same SFASW media. Ground electrode was placed in the SFSW of the dish and potential was recorded between the ground and suction electrode. Amplifier was additionally grounded to reduce electronic noise. Events were amplified using Grass CP511 Amplifiers and recorded through WinWCP software at a sampling interval of 0.1-0.5 ms at 1000x amplification, with a 1 kHz high pass and 3 Hz low pass filter.

Recordings were analyzed by filtering with a low-pass Butterworth filter, extracting events lasting an average of 20-40 ms with peaks greater than 3 standard deviations above the mean [56]. All extracted events were visualized and incorrectly identified waves were dropped from analysis. Events were clustered with Principal Components Analysis and a *k*-means clustering. Features utilized are described in Supplemental Figure 2A. Clustered events were then averaged together to visualize characteristic waveforms.

## Supplemental Figures

**Figure 1 Supplemental.**
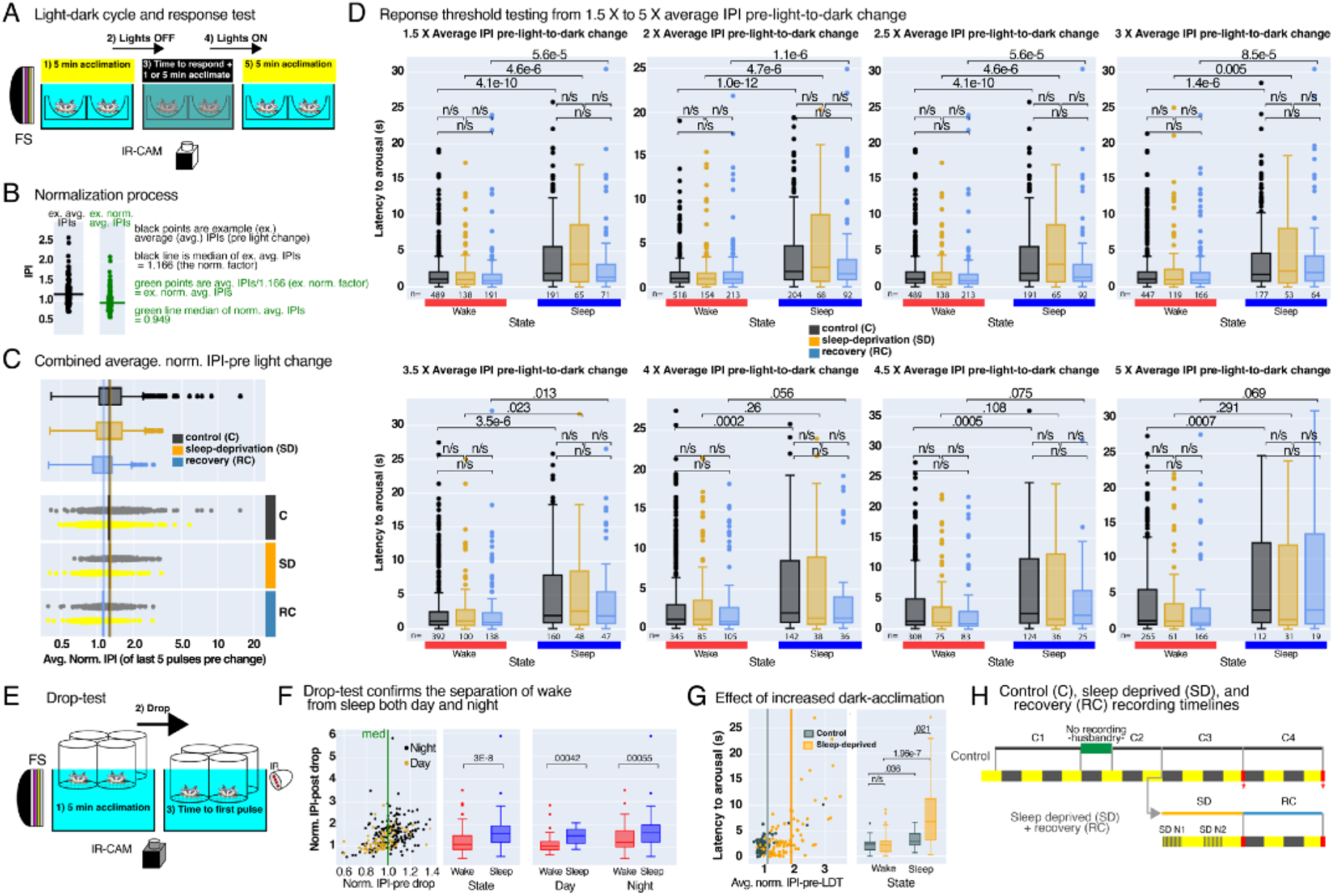
LTA paradigm, normalization, response threshold testing, Drop-test based LTA, and longterm recording timeline. **(A)** We adapted the previously developed drop-test [7], to make infrared recordings. **(B)** The average of each 5-IPI set is calculated generating the light- and dark-average IPIs, left panel in black is example for one jellyfish. Each measurement is divided by its respective **normalization factor** for a **dimensionless normalized IPI**, to generate the **normalized avg-IPI**, right panel in green. **(C)** A threshold is required to distinguish sleep from wake, we used a **median threshold** (the combined normalized average of light- and dark-IPIs pre-LDT,) for control, SD, and RC animals. **(D)** To measure the LTA a **response time** (time from the light-dark switch until pause response) is required. We tested various thresholds for defining a pause, from 1.5X to 5X the avg-IPI pre-light-dark change, all showed segregation of sleep from wake. p-values calculated using one-way ANOVA, followed by Wilcox rank sum test. **(E)** *Cassiopea* were placed in a 2×3 array of clear screen-bottom cylinders in a temperature controlled pulse tracking setup. Every 10min for 3 hours the animals are “dropped” to determine their LTA (n=12, 155 events). Drop-tests were performed in the light and in the dark. **(F)** IPI pre-drop is compared to the IPI post-drop. Taking the median (med, green line) of pre-drop IPIs generates a thresh-old for comparing post-drop IPIs (left panel). Box-plots compare the below median wake (red) and above median sleep (blue) responses (righty panel), below median (wake-red) and above median (blue-sleep) occur both day and night. **(G)** To measure the LTA of animals with greater dark acclimation, LTA was measured in control and sleep-deprived animals. n = 21 BL and n =16 SD, 150 BL and 90 SD events, p-values calculated using one-way ANOVA, followed by Wilcox rank sum test. **H)** The behavioral recording paradigm includes an initial control recording (C1), that includes 2 full nights and 2 full days, followed by a husbandry period for feeding and cleaning. Jellies are then returned to the recording setup for an additional night and day of recording (C2). After this point, jellies are either going to receive two more recordings, each with 2 nights and 2 days of control, or 2 nights of sleep deprivation (SD), followed by 2 nights and days of recovery (RC).

**Figure 2 Supplemental.**
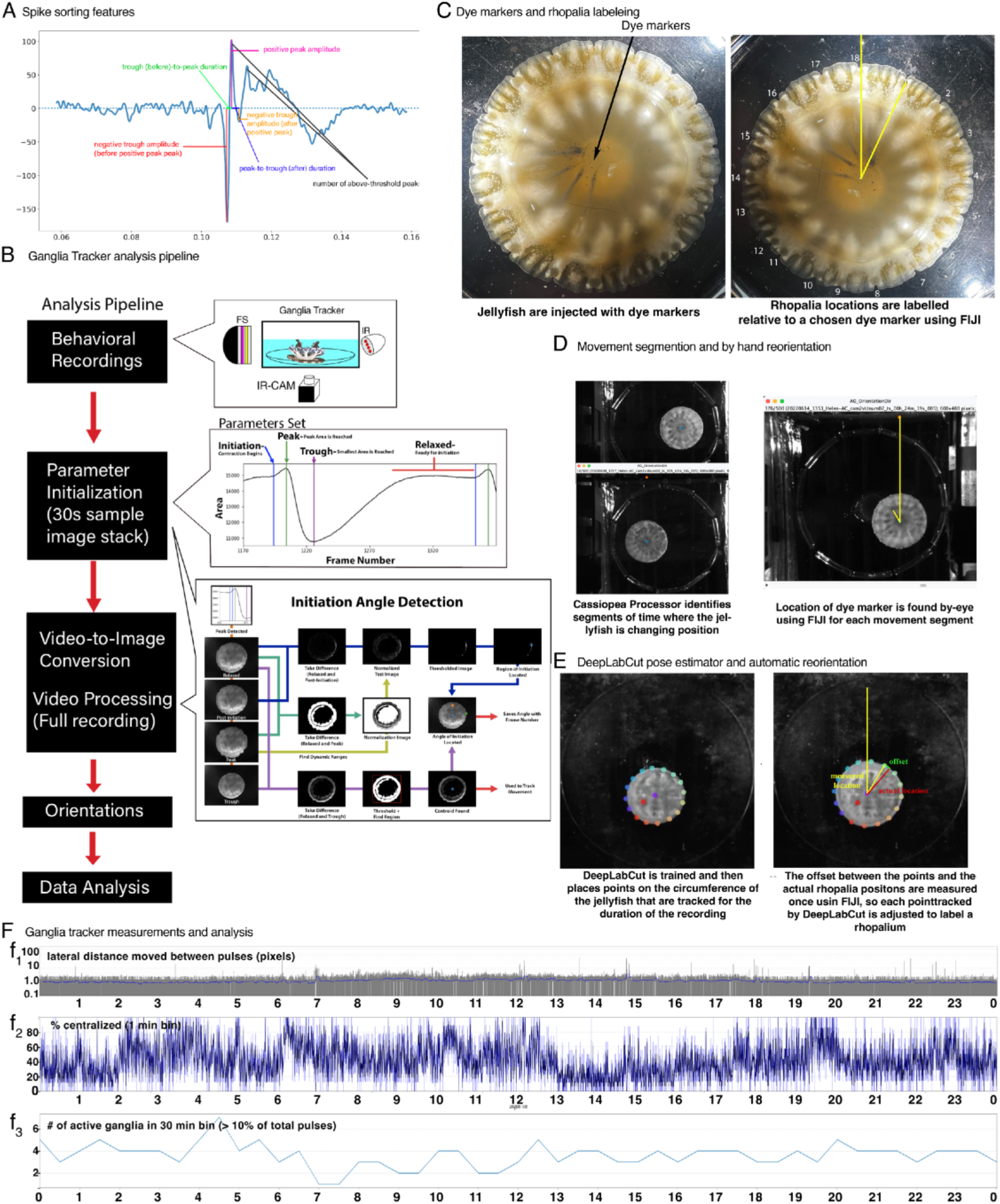
*Cassiopea* orientation, analysis pipeline, and lateral distance moved, centralization, and active ganglia count metrics. **(A)** Sample waveform with features used for spike sorting through PCA and k-means clustering. Features chosen describe the amplitude of troughs and peaks, duration from each trough to peak, and the number of peaks above a set threshold. **(B)** Steps of CassiopeaProcessor pipeline, from long-term behavioral recordings to final acquisition of data on initiation angles. Visualization of key steps on the right, including the behavioral recording setup, the initialization parameters that are manually set, and the method of initiation angle detection through thresholding of frames identified by using initialization parameters. **(C)** India ink (See Methods) injections aline with rhopalia are visible as dark lines in infrared. The dye mark that is clearest after several days of recording is then used to assign rhopalia numbers and measure the angles of each. **(D)** When animals move 20 pixels between contractions they are considered moving, and the video processing stops analysis and once the jelly is not moving for 30 seconds it begins again. The dye marks allow for re-orientation of the animals by-eye. **(E)** DeepLabCut pose estimator has been trained to track points on the bell margin, which are then adjusted to mark rhopalia. Reorientation is then accomplished automatically. **(F)** Ganglia Tracker metrics. **(f**_**1**_) The centroid of the jellyfish is determined every pulse and the distance in pixels of the centroid between pulses is measured. **(f**_**2**_) The frequency of consecutive pulses from a single ganglion is plotted as metric for network centralization, binned every 1 min. **(f**_**3**_**) T**he number of active (above 10% of pulses) ganglia in 30 min bins.

**Figure 3 Supplemental.**
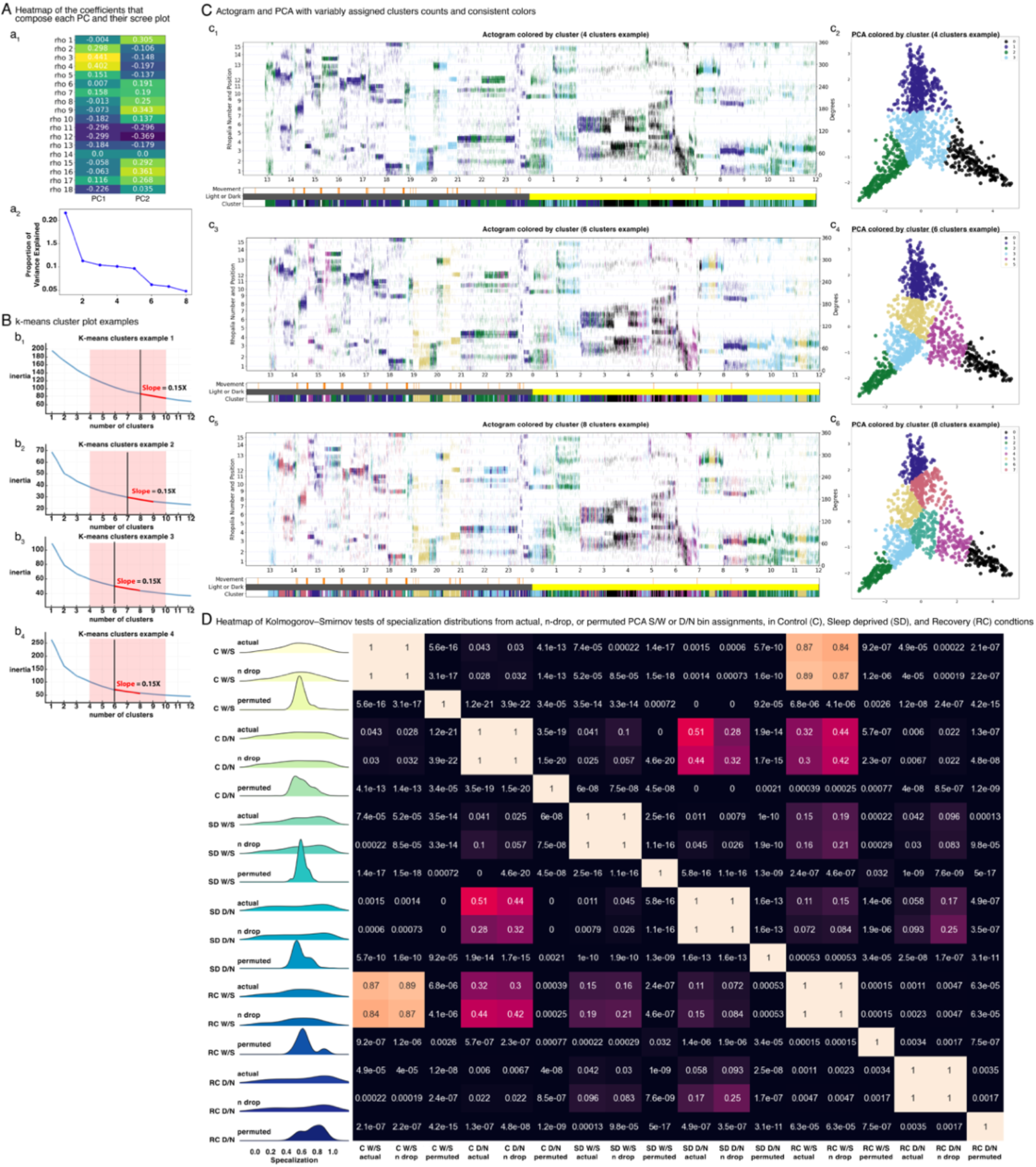
Assessment of clustering model, k-means cluster plots, and sleep vs. wake and night vs. day specialization comparisons using actual, n-drop or permuted cluster data. **(A)** Heatmap of the coefficients of each feature in the linear combination that composes each PC and associated scree plot. **(a**_**1**_) The absolute value is indicative of the importance of each feature in capturing the variance across the original dataset. **(a**_**2**_) Sample scree plot describing the proportion of explained variance captured by each additional principal component. The first PC captures about 23% of total variance, while the 2nd PC captures approximately 12% of the total variance. Current model using 2 PCs encompasses 35% of total variance. **(B)** K-means algorithm is an unsupervised machine learning algorithm. Each 24-hour recording is divided into K number of clusters. Every cluster has its centroid which is calculated by averaging the data points of that cluster. Inertia is the measure of intra-cluster distances, or how far away the datapoint is from the cluster’s centroid. **(b**_**1-4**_), Empirically, eight clusters are near or below the elbow (black line) of every each 24-hour recording inertia plot, where the red line indicates the approximate slope of the elbow region, associated with ∼8 clusters. **(C)** Actograms from a single animal but assigning **(c**_**1-2**_) 4, **(c**_**3-4**_) 6, or **(c**_**5-6**_) 8 clusters to visually compare actogram clustering accuracy. The actograms **(c**_**1**,**3**,**5**_) were then re-colored using the cluster color associated with each 1-min bin for their associated PCA clustering **(c**_**2**,**4**,**6**_**). (D)** PCA sleep/wake and night/day specialization analysis of n = 14 control animals with 385 clusters, n = 14 SD animals with 223 clusters, and n=5 RC animals and 54 clusters. Ridge plots compare the distributions the specialization % for each cluster, separated by condition, visualized using matplotlib gaussian kernel density estimator. Y-axis is normalized density, and X-axis is the frequency of % sleep (W/S), or % night (N/D), per cluster; 1 is maximum sleep or night specialization, 0 is maximum wake or day specialization, and 0.5 is unspecialized. Distribution names are: Control (C), sleep deprived (SD), and recovery (RC), using the experimentally collected data (actual), two bins dropped per cluster (n drop), or permuted bin assignments (permuted). Heat map p-values determined by Kolmogorov-Smirnov test.

**Figure 4 Supplemental.**
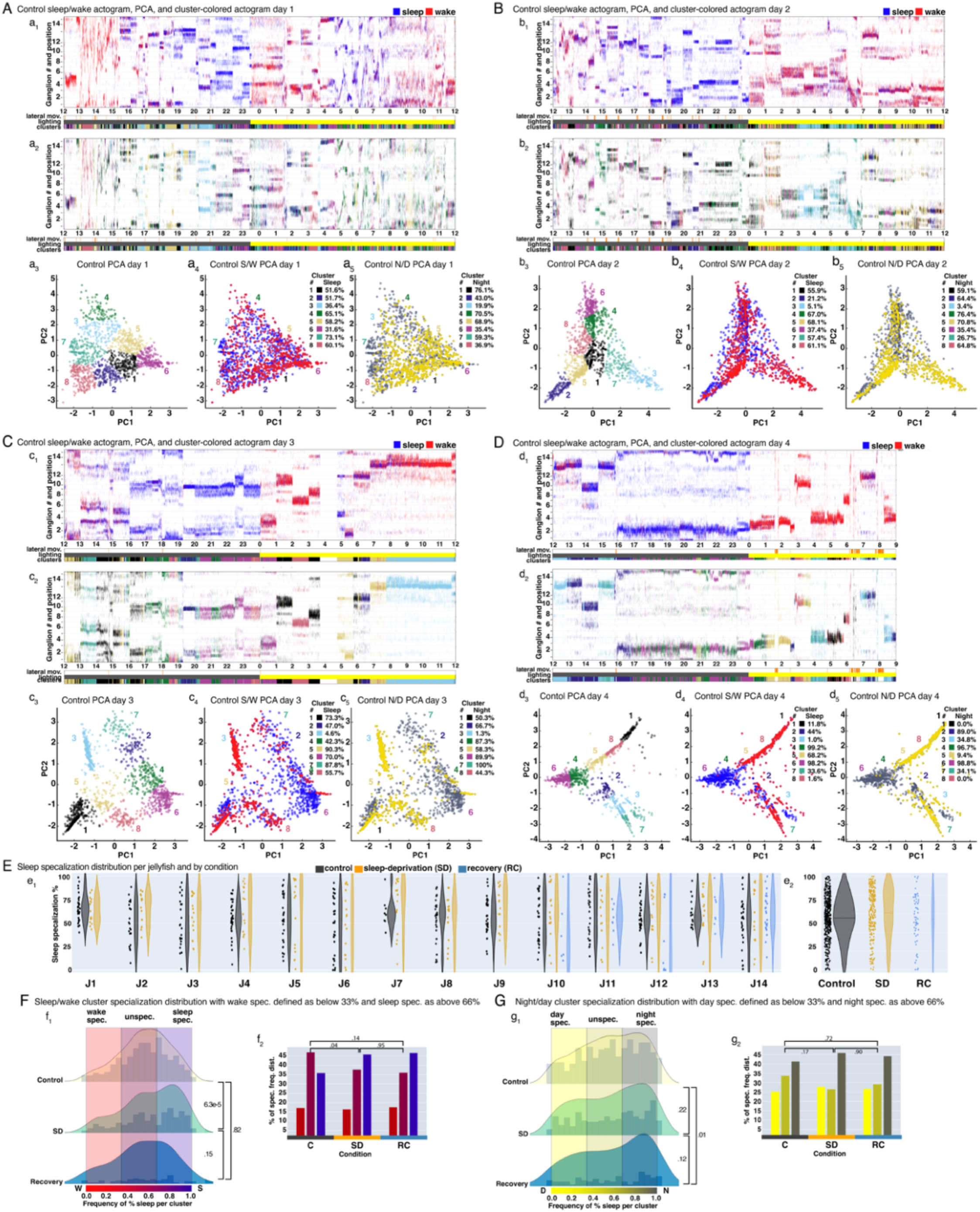
Distribution of sleep specialization, PCA timeline and associated actograms, sleep specialization across all 14 tracked animals, and assessment of the maximum range for specialization and ganglia change. **(A-D)** Actograms and PCAs from a single animal four consecutive days and nights, using ganglia activity 1-min bins, and k-means clustering. **(a**_**1**_, **b**_**1**_, **c**_**1**_, **d**_**1**_), Consecutive actograms, **(a**_**2**_, **b**_**2**_, **c**_**2**_, **d**_**2**_) recolored actograms based on PCA cluster data in **(a**_**3**_, **b**_**3**_, **c**_**3**_, **d**_**3**_). PCA bins recolored based on **(a**_**4**_, **b**_**4**_, **c**_**4**_, **d**_**4**_) sleep/wake and **(a**_**5**_, **b**_**5**_, **c**_**5**_, **d**_**5**_) day/night bin assignments. Specialization per cluster is then calculated as the ratio of each cluster. **(E)** Sleep specialization distribution and KDE for **(e**_**1**_) each analyzed jellyfish, or per condition **(e**_**2**_). n = 14 control animals with 385 clusters, n = 14 SD animals with 223 clusters, and n=5 RC animals and 54 clusters. Visualized using matplotlib Gaussian kernel density estimator. Y-axis is the frequency of % sleep per cluster; 1 is maximum sleep specialized, 0 is maximum wake specialized, and 0.5 is unspecialized. **(A-G)** Same visualization of specialization distribution as in Figure 5 D-E, except the regions considered specialized are sleep **(d**_**2**_) or night **(e**_**2**_) (0.66 - 1.0), **(d**_**2**,_ **e**_**2**_) unspecialized (0.33-0.66), and wake **(d**_**2**_) or day **(e**_**2**_) specialized (0.0 - 0.33). p-values determined by 2×3 chi-square.

## References

1. Tononi, G. and Cirelli, C., 2006. Sleep function and synaptic homeostasis. Sleep medicine reviews, 10(1), pp.49–62.

2. Shaw, P.J., Cirelli, C., Greenspan, R.J. and Tononi, G., 2000. Correlates of sleep and waking in Drosophila melanogaster. Science, 287(5459), pp.1834–1837.

3. Eugene, A.R. and Masiak, J., 2015. The neuroprotective aspects of sleep. MEDtube science, 3(1), p.35. Joiner, W.J. (2016). Unraveling the evolutionary determinants of sleep. Curr. Biol. 26, R1073–R1087.

4. Walker, M.P. and Stickgold, R., 2004. Sleep-dependent learning and memory consolidation. Neuron, 44(1), pp.121–133.

5. Tononi, G. and Cirelli, C., 2014. Sleep and the price of plasticity: from synaptic and cellular homeostasis to memory consolidation and integration. Neuron, 81(1), pp.12–34.

6. Frank, M.G. and Cantera, R., 2014. Sleep, clocks, and synaptic plasticity. Trends in neurosciences, 37(9), pp.491–501.

7. Nath, R.D., Bedbrook, C.N., Abrams, M.J., Basinger, T., Bois, J.S., Prober, D.A., Sternberg, P.W., Gradinaru, V. and Goentoro, L., 2017. The jellyfish Cassiopea exhibits a sleep-like state. Current Biology, 27(19), pp.2984–2990.

8. Kanaya, H.J., Park, S., Kim, J.H., Kusumi, J., Krenenou, S., Sawatari, E., Sato, A., Lee, J., Bang, H., Kobayakawa, Y. and Lim, C., 2020. A sleep-like state in Hydra unravels conserved sleep mechanisms during the evolutionary development of the central nervous system. Science advances, 6(41), p.eabb9415.

9. Bosch, T.C., Klimovich, A., Domazet-Lošo, T., Gründer, S., Holstein, T.W., Jékely, G., Miller, D.J., Murillo-Rincon, A.P., Rentzsch, F., Richards, G.S. and Schröder, K., 2017. Back to the basics: cnidarians start to fire. Trends in Neurosciences, 40(2), pp.92–105. Joiner, W.J., 2016. Unraveling the evolutionary determinants of sleep. Current biology, 26(20), pp.R1073–R1087.

10. Campbell, S.S. and Tobler, I., 1984. Animal sleep: a review of sleep duration across phylogeny. Neuroscience & Biobehavioral Reviews, 8(3), pp.269–300.

11. Greenspan, R.J., Tononi, G., Cirelli, C. and Shaw, P.J., 2001. Sleep and the fruit fly. Trends in neurosciences, 24(3), pp.142–145.

12. Zhdanova, I.V., Wang, S.Y., Leclair, O.U. and Danilova, N.P., 2001. Melatonin promotes sleep-like state in zebrafish. Brain research,903(1-2), pp.263–268.

13. Singh, K., Chao, M.Y., Somers, G.A., Komatsu, H., Corkins, M.E., Larkins-Ford, J., Tucey, T., Dionne, H.M., Walsh, M.B., Beaumont, E.K. and Hart, D.P., 2011. C. elegans Notch signaling regulates adult chemosensory response and larval molting quiescence. Current Biology, 21(10), pp.825–834.

14. Singh, K., Ju, J.Y., Walsh, M.B., DiIorio, M.A. and Hart, A.C., 2014. Deep conservation of genes required for both Drosophila melanogaster and Caenorhabditis elegans sleep includes a role for dopaminergic signaling. sleep, 37(9), pp.1439–1451.

15. Roelofs, K., 2017. Freeze for action: neurobiological mechanisms in animal and human freezing. Philosophical Transactions of the Royal Society B: Biological Sciences, 372(1718), p.20160206.

16. Anafi, R.C., Kayser, M.S. and Raizen, D.M., 2019. Exploring phylogeny to find the function of sleep. Nature Reviews Neuroscience, 20(2), pp.109–116.

17. Achermann, P., Borbély, A.A., Kryger, M.H., Roth, T. and Dement, W.C., 2017. Sleep homeostasis and models of sleep regulation.

18. Allada, R. and Siegel, J.M., 2008. Unearthing the phylogenetic roots of sleep. Current biology, 18(15), pp.R670–R679.

19. Machado, R.B., Suchecki, D. and Tufik, S., 2005. Sleep homeostasis in rats assessed by a long-term intermittent paradoxical sleep deprivation protocol. Behavioural brain research, 160(2), pp.356–364.

20. Passano, L.M., 1982. Scyphozoa and cubozoa. Electrical conduction and behaviour in ‘simple’invertebrates. Oxford University Press, New York, pp.149–202.

21. Schwab, W.E., 1977. The ontogeny of swimming behavior in the scyphozoan, Aurelia aurita. I. Electrophysiological analysis. The Biological Bulletin, 152(2), pp.233–250.

22. Passano, L.M., 1965. Pacemakers and activity patterns in medusae: homage to Romanes. American zoologist, 5(3), pp.465–481.

23. Passano, L.M., 1973. Behavioral control systems in medusae; a comparison between hydro-and dscyphomedusae. Publications of the Seto Marine Biological Laboratory,20, pp.615–645.

24. Nakanishi, N., Hartenstein, V. and Jacobs, D.K., 2009. Development of the rhopalial nervous system in Aurelia sp. 1 (Cnidaria, Scyphozoa). Development genes and evolution, 219(6), pp.301–317.

25. Martin, V.J., 1992. Characterization of a RFamide-positive subset of ganglionic cells in the hydrozoan planular nerve net. Cell and tissue research, 269(3), pp.431–438.

26. Takahashi, T. and Takeda, N., 2015. Insight into the molecular and functional diversity of cnidarian neuropeptides. International Journal of Molecular Sciences, 16(2), pp.2610–2625.

27. Findeisen, M., Rathmann, D. and Beck-Sickinger, A.G., 2011. RFamide peptides: structure, function, mechanisms and pharmaceutical potential. Pharmaceuticals, 4(9), pp.1248–1280.

28. Satterlie, R.A. and Eichinger, J.M., 2014. Organization of the ectodermal nervous structures in jellyfish: scyphomedusae. The Biological Bulletin, 226(1), pp.29–40.

29. Stöckl, A.L., Petie, R. and Nilsson, D.E., 2011. Setting the pace: new insights into central pattern generator interactions in box jellyfish swimming. PloS one, 6(11), p.e27201.

30. Mackie, G.O., 2004. Central neural circuitry in the jellyfish Aglantha. Neurosignals,13(1-2), pp.5–19.

31. Pallasdies, F., Goedeke, S., Braun, W. and Memmesheimer, R.M., 2019. From single neurons to behavior in the jellyfish Aurelia aurita. Elife,8, p.e50084.

32. Kanwisher, N., 2010. Functional specificity in the human brain: a window into the functional architecture of the mind. Proceedings of the National Academy of Sciences, 107(25), pp.11163–11170.

33. Saper, C.B., Chou, T.C. and Scammell, T.E., 2001. The sleep switch: hypothalamic control of sleep and wakefulness. Trends in neurosciences, 24(12), pp.726–731.

34. Rensing, L. and Ruoff, P., 2002. Temperature effect on entrainment, phase shifting, and amplitude of circadian clocks and its molecular bases. Chronobiology international, 19(5), pp.807–864.

35. Kavanau, J.L., 1971. Locomotion and activity phasing of some medium-sized mammals. Journal of Mammalogy, 52(2), pp.386–403.

36. Cao, W. and Edery, I., 2017. Mid-day siesta in natural populations of D. melanogaster from Africa exhibits an altitudinal cline and is regulated by splicing of a thermosensitive intron in the period clock gene. BMC evolutionary biology, 17(1), pp.1–17.

37. Abdi, H. and Williams, L.J., 2010. Principal component analysis. Wiley interdisciplinary reviews: computational statistics, 2(4), pp.433–459.

38. Hartigan, J.A. and Wong, M.A., 1979. Algorithm AS 136: A k-means clustering algorithm. Journal of the royal statistical society. series c (applied statistics), 28(1), pp.100–108.

39. Weissbourd, B., Momose, T., Nair, A., Kennedy, A., Hunt, B. and Anderson, D.J., 2021. A genetically tractable jellyfish model for systems and evolutionary neuroscience. Cell, 184(24), pp.5854–5868.

40. Szymanski, J.R. and Yuste, R., 2019. Mapping the whole-body muscle activity of Hydra vulgaris. Current Biology, 29(11), pp.1807–1817.

41. Saper, C.B., Fuller, P.M., Pedersen, N.P., Lu, J. and Scammell, T.E., 2010. Sleep state switching. Neuron, 68(6), pp.1023–1042.

42. Nichols, A.L., Eichler, T., Latham, R. and Zimmer, M., 2017. A global brain state underlies C. elegans sleep behavior. Science, 356(6344), p.eaam6851.

43. Zielinski, M.R., McKenna, J.T. and McCarley, R.W., 2016. Functions and mechanisms of sleep. AIMS neuroscience, 3(1), p.67.

44. Mignot, E., 2008. Why we sleep: the temporal organization of recovery. PLoS biology, 6(4), p.e106.

45. Klimovich, A., Giacomello, S., Björklund, Å., Faure, L., Kaucka, M., Giez, C., Murillo-Rincon, A.P., Matt, A.S., Wil-loweit-Ohl, D., Crupi, G. and de Anda, J., 2020. Prototypical pacemaker neurons interact with the resident microbiota. Proceedings of the National Academy of Sciences, 117(30), pp.17854–17863.

46. Martinez, D., Santin, J.M., Schulz, D. and Nadim, F., 2019. The differential contribution of pacemaker neurons to synaptic transmission in the pyloric network of the Jonah crab, Cancer borealis. Journal of neurophysiology, 122(4), pp.1623–1633.

47. Song, J., Pallucchi, I., Ausborn, J., Ampatzis, K., Bertuzzi, M., Fontanel, P., Picton, L.D. and El Manira, A., 2020. Multiple rhythm-generating circuits act in tandem with pacemaker properties to control the start and speed of locomotion. Neuron, 105(6), pp.1048–1061.

48. Blankenship, A.G. and Feller, M.B., 2010. Mechanisms underlying spontaneous patterned activity in developing neural circuits. Nature Reviews Neuroscience, 11(1), pp.18–29.

49. Homma, S., Kobayashi, Y., Kosugi, S., Ohashi, M., Kanda, T., Okamoto, H. and Hatakeyama, K., 2008. Postoperative reorganization of gastric pacemaker activity in patients after an extended period following distal gastrectomy. Journal of Smooth Muscle Research,44(3+ 4), pp.113–122.

50. Torrente, A.G., Zhang, R., Zaini, A., Giani, J.F., Kang, J., Lamp, S.T., Philipson, K.D. and Goldhaber, J.I., 2015. Burst pacemaker activity of the sinoatrial node in sodium–calcium exchanger knockout mice. Proceedings of the National Academy of Sciences, 112(31), pp.9769–9774.

51. Dobs, K., Martinez, J., Kell, A.J. and Kanwisher, N., 2022. Brain-like functional specialization emerges spontaneously in deep neural networks. Science advances, 8(11), p.eabl8913.

52. Lewis Ames, C., 2018. Medusa: a review of an ancient cnidarian body form. Marine organisms as model systems in biology and medicine, pp.105–136.

53. Mascetti, G.G., 2016. Unihemispheric sleep and asymmetrical sleep: behavioral, neurophysiological, and functional perspectives. Nature and Science of Sleep,8, p.221.

54. Vyazovskiy, V.V., Olcese, U., Hanlon, E.C., Nir, Y., Cirelli, C. and Tononi, G., 2011. Local sleep in awake rats. Nature, 472(7344), pp.443–447.

55. Mathis, A., Mamidanna, P., Cury, K.M., Abe, T., Murthy, V.N., Mathis, M.W. and Bethge, M., 2018. DeepLabCut: markerless pose estimation of user-defined body parts with deep learning. Nature neuroscience, 21(9), pp.1281–1289.

56. Laboy-Juárez, K.J., Ahn, S. and Feldman, D.E., 2019. A normalized template matching method for improving spike detection in extracellular voltage recordings. Scientific reports, 9(1), pp.1–12.

